# The impact of nerve injury on the immune system across the lifespan is sexually dimorphic

**DOI:** 10.1101/2025.04.24.650472

**Authors:** Wen Bo Sam Zhou, Xiang Qun Shi, Alain P Zhang, Magali Millecamps, Jeffrey S. Mogil, Ji Zhang

**Author notes:** Corresponding Author Dr. Ji Zhang, The Alan Edwards Centre for Research on Pain McGill University, Lyman Duff Medical Building, Suite 203 3775 University Street, Montreal, H3A 2B4, QC, Canada.

## Abstract

Although nerve injury-associated neuroinflammation contributes to neuropathic pain, the long-term impact of such injury on systemic homeostasis and its potential role in pain remains elusive. In this study, we aim to understand the systemic changes that are present alongside chronic pain in nerve-injured male and female mice across their lifespan. We monitored mechanical and cold sensitivity in male and female mice starting at the age of 3–4 months old when they received spared nerve injury (SNI), up to 20-month post-injury. Alongside, we collected blood samples to track changes in immune cells with flow cytometry, and to assess inflammation-related serum proteome using a 111-target Proteome Profiler. We also transferred serum from sham/SNI mice to naïve mice to determine the potential of systemic contribution to pain. While nerve injury did not affect immune cell composition in the blood, it triggered a long-lasting disturbance of molecular profile in the serum of sham/SNI mice, in a sex-dependent manner. Compared to sham surgery, nerve injury amplified regulation of inflammatory proteins in males, but slightly reduced it in females. These changes in the serum occurred in parallel with long-lasting mechanical and cold hypersensitivity in the nerve-injured mice. Both male and female SNI serum induced hypersensitivity when transferred to naïve mice, regardless of a sex-matched or sex-crossed transfer. Our results highlight that a local nerve injury can have persistent systemic impact. Injury-associated systemic inflammation could contribute to neuropathic pain, but the underlying mechanisms may be sexually dimorphic.

## Introduction

Peripheral nerve injury has an annual incidence between 7–17 per 100,000 persons around the world (1, 2). It affects both motor and sensory functions, leading to disability and poor quality of life (3). Unlike those in the central nervous system, neurons in the peripheral nervous system can regenerate after damage. However, the success of the regeneration and functional recovery will depend on the nature and the severity of injury—stretch, laceration, compression, transection or avulsion—and the changes in local and systemic microenvironment. Nerve injury not only triggers inflammation at the site of damage, but also remotely in the dorsal root ganglia (DRG) (4) and in the spinal cord (5). Such inflammatory reactions within nervous system are mediated by both resident and infiltrating immune cells, along with inflammatory mediators that play crucial roles in Wallerian degeneration and subsequent regeneration processes (6).

In patients, direct evidence of nerve injury-associated neuroinflammation is limited, partly due to the invasive nature of accessing nervous tissues. However, increasing evidence points to an increase of various proinflammatory cytokines and chemokines in the serum of neuropathy patients, especially among those who experience neuropathic pain (7, 8). This suggests the potential impact of a nerve injury on the immune system, although the direct correlation between the levels of circulating proinflammatory mediators and injury-associated painful neuropathy remains controversial (9). Several preclinical studies have also provided similar evidence using different animal models (10, 11) to endorse the clinical observations that a local nerve injury may have a systemic impact. However, as the vast majority of preclinical research on nerve injury and chronic pain has been conducted on 2–3 month-old male rodents, and terminated before or at 1 month following injury (12), the long-term impact of a simple peripheral nerve injury on the immune system, and the potential of such immune disturbance in contributing to long-lasting injury-triggered pain, remains obscure.

Furthermore, although men are at least twice as likely to suffer from nerve injury than women (13, 14), women are far more likely to develop neuropathic pain (15, 16). Preclinical studies show mixed findings on the sex difference in nerve regeneration following damage on the nerve (17, 18). Since the immune system in both humans and in rodents displays sexually dimorphic signatures (19), it is reasonable to hypothesize that the male and female immune system may respond differently to an injury. Further dissection of the sex difference of injury-triggered systemic inflammation could advance our understanding on the role of inflammation in nerve degeneration/regeneration and the development of chronic pain.

In this study, we aim to characterize the systemic inflammation developed following a non-resolving peripheral nerve injury, and the potential of such inflammation in the development of neuropathic pain. By using spared sciatic nerve injury (SNI) as a model, we assessed the number of circulating immune cells and serum proteome of sham- and SNI-operated mice of both sexes across the lifespan from 1–20 months post-injury. Our data reveal that nerve injury has a long-term impact on the immune system and generates persistent systemic chronic inflammation, which exhibits clear sexually dimorphic features. Transferring serum from either male or female SNI mice to naïve mice induced mechanical and cold allodynia. This suggests that nerve injury-associated systemic chronic inflammation contributes to pain hypersensitivity, but could be mediated by different molecular mechanisms in male and female mice.

## 2. Materials and Methods

### 2.1 Animals

Male and female 3–23-month-old C57BL/6J mice, bred in house from breeders obtained from The Jackson Laboratory (Bar Harbor, ME), were included in the study. Mice were housed 2–5 per cage in a temperature- and humidity-controlled environment with a 12/12-hour light/dark cycle, with the light phase beginning at 07:00, and were given access to rodent chow (Teklad, #2920) and tap water *ad libitum*. All protocols were approved by the McGill University Animal Care and Use Committee (protocol# 5775) and conducted according to the guidelines of the Canadian Council on Animal Care.

### 2.2 Nerve Injury Model

The spared nerve injury (SNI) was performed under isoflurane/oxygen anesthesia as originally described in rats by Decosterd and Woolf (20). Briefly, the sciatic nerve, along with the sural, tibial, and common peroneal branches was exposed. The tibial and common peroneal nerves were transected and tightly ligated with 7.0-silk threads, while the sural nerve was spared. For the sham surgeries, the sciatic nerve branches were exposed but not transected. Forty mice at the age of 2-3 months were used for the nerve injury model, with 10 mice per surgery per sex.

Mice from each cage were randomly selected for sham or SNI surgery and were co-housed in the same cages. Additional groups of male and female young (3-4 months old) naïve mice were included for immune cells and serum analysis (n=11/group) and for serum transfer experiments (n=5/condition).

### 2.3 Behavioral Analysis

Behavioral experiments were conducted between 09:00 and 15:00. Mice were habituated to the testing environment for at least 1 hour before the start of testing. For measurements of mechanical sensitivity on the paw, withdrawal thresholds were assessed with calibrated von Frey filaments (Stoelting, Wood Dale, IL) using the up-down method of Dixon (21). Mice were placed on a metal mesh floor (1/8 x 1/8”) within small Plexiglas cubicles (9 x 5 x 5-cm high). A set of calibrated von Frey filaments (ranging from 0.16–1.40-g bending force) were applied to lateral hind paw region innervated by the sural nerve until they bent. A decrease in paw-withdrawal thresholds indicates mechanical hypersensitivity.

Cold sensitivity was evaluated with the acetone test. A drop of acetone, approximately 25 µL in volume, was applied to the lateral surface of the hind paw. The duration of acetone-evoked nocifensive behaviors (i.e., flinching, licking, or lifting) after the application of acetone within a 1-minute observation period was recorded. An increase in the duration of nocifensive behaviors indicates cold hypersensitivity.

In experiments with SNI- and sham-operated mice, neuropathic pain behavior was monitored with the two aforementioned tests at 1, 7, 10, 12, 15, 18 and 20 months post-injury. The investigator performing behavioral experiments was blinded to the groups and treatment, although blinding to sex was not possible. In experiments with naïve mice that received an intravenous transfer of serum from sham or SNI mice, behavioral tests were performed on the walking pad of both left and right paws, and the results from both sides were averaged. Pain behavior was assessed once every 2 days starting at day 1 following serum administration, and until mechanical and cold sensitivity return to their baseline levels. An additional test was performed the day after return to confirm the full recovery of the pain episode.

### 2.4 Serum Collection and Transfer

Serum was sampled via the submandibular vein from mice at 1, 6, 12, 15, and 20 months following SNI or sham surgery. Serum from young (3-4 months old) naïve male and female mice were also collected for analysis. Blood was harvested in Microvette 200 serum collection tubes (ref. 20.1291, Sarstedt) and allowed to clot for 30 minutes at room temperature before centrifugation at 10,000 x *g* for 5 minutes to isolate the serum. To avoid potential changes due to circadian rhythms, all blood samples were collected between 10:00-12:00. The serum was stored at −80 C° until use.

Transfer of serum from SNI or sham mice to naïve mice were carried out via tail vein injections using 1-mL syringes and 28-G needles. Each intravenous tail vein injection consisted of 100 µL serum pooled from 5–10 SNI or sham mice.

### 2.5 Proteome Profiler Assay and blot image analysis

For protein analysis, serum from SNI or sham mice at each time point post-injury were pooled to 200 µL, using equal volumes of serum per mouse, from 3–10 mice in each condition. Each pooled sample was applied to an ELISA membrane from the Proteome Profiler Mouse XL Cytokine Array Kit (ARY028, R&D Systems) and developed according to the manufacturer’s instructions. The sample was incubated with the membrane for 40 hours. The protein signals on the membrane were detected using an Amersham™ Imager 600. Quantification of the signals was performed manually in ImageJ using the Protein Array Analyzer plugin by placing an array template on the image. The result for each protein signal was averaged from two manual quantifications of the duplicate spots. After subtracting the background from each signal, each data point was processed as a ratio and log2 fold change, or as a difference between two conditions for subsequent plotting.

To test for potential correlations with each factor (sex, nerve injury, age), data subsets were created with only one varying factor. For example, to correlate with age, the “male_sni” subset includes data for each age level including male SNI 1 month, male SNI 12 month, and male SNI 20 months. Each subset was set as an input into a two-stage correlation categorization function. The first stage calculated the relative percent change between subsequent factor levels for each analyte. For a potential correlation to exist, the magnitude of the change between at least one pair of subsequent levels had to meet or exceed 100% or the analyte would be categorized as uncorrelated with the factor for the input data subset.

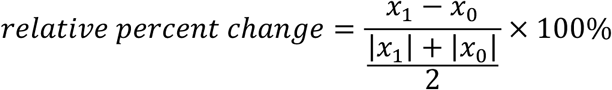

Correlations were either categorized as “positive” or “negative” demonstrated for an increasing or decreasing trend in analyte quantity with factor. For correlations with age, a third option of “no correlation” was also possible for unordered analyte quantities since there were three age levels tested.

To determine if the correlations persisted across other factors, the correlation categorization function was applied for all subsets. For the age correlation example, the subsets *male_sham, male_sni, female_sham, and female_sni* were each inputted into the correlation categorization function to determine if an age correlation was indicated regardless of sex or treatment. The correlation categorization results were then cross-referenced and tallied to produce a match score represented as the proportion of subsets that which showed the same trend for an analyte, with a 100% match score indicating that the correlation for a particular analyte was indicated for all combinations of factors and levels.

Data from ImageJ was imported into a Jupyter notebook using Python and reformatted with the Pandas library. Calculations for correlation indication were performed with the NumPy library. The correlation plots were generated with the Matplotlib library using seaborn-like styling.

### 2.7 Luminex

Individual serum samples from naïve mice and from mice at 6 and 15 months following SNI or sham surgery were used to validate regulation pattern observed with pooled samples in Proteome Profiler analysis. Targets were randomly selected based on commercial kit availability and overlap in the Proteome Profiler assay, which include G-CSF (CSF-3), MCP-1 (CCL-2), MIP-1 beta (CCL-4), and TNF alpha (ThermoFisher). The assay was performed according to manufacturer’s protocol with an overnight incubation and acquired with the MAGPIX instrument. The Luminex xPONENT software (version 4.2.1705.0) was used to determine cytokine concentrations using a 5-parameter logistic regression on 8 standard curve dilutions for each cytokine.

### 2.8 Flow cytometry

Blood immune cells were analyzed at 1, 7, 12, 15, and 18 months post-injury. Blood was sampled via the submandibular vein; 50 μL was collected and transferred into tubes with Alsever’s solution to prevent coagulation. To avoid potential changes due to circadian rhythms, all blood samples were collected between 10:00 am-12:00 pm. Red blood cells were lysed with two rounds of incubation with ACK (ammonium-chloride-potassium) lysing buffer for 5 minutes on ice to remove erythrocytes. Samples were blocked with 2.4G2 Fc block for 30 minutes prior to 30 minutes of antibody staining, both procedures were conducted on ice. Cells were labelled with two antibody panels in the two separate tubes: Panel 1: allophycocyanin (APC) anti-CD115 (135510, BioLegend) and PerCP-Cy5.5 anti-CD11b (101228, BioLegend) (monocytes), APC-Cy7 anti-Ly6G (560600, BD Biosciences) (neutrophils), and fluorescein isothiocyanate (FITC) anti-NK1.1 (108718, BioLegend) (NK cells); Panel 2: PE anti-CD4 (553048, BD Biosciences) and APC anti-CD8a (100712, BioLegend) (CD4 and CD8 T cells), and FITC anti-CD19 (115506, BioLegend) (B cells). All antibodies were diluted 1:50 in blocking buffer.

The PE anti-CD3e antibody (100308, BioLegend) was only used at the 7 months post-injury time point, where the anti-CD4, anti-CD8, and anti-CD19 antibodies were not used. The gating strategies for each specific cell type are shown in Spplementary Fig S1. Flow cytometry acquisition was performed on the BD FACSCanto II machine using the BD FACSDiva software. Total leukocyte numbers were calculated based on the gated value in the FSC and SSC scatter and the absolute cell count was determined using Precision Count beads (424902, BioLegend). The population of “Other Granulocytes” was gated based on Ly6G-negative, SSC-high cells. Data was analyzed with FlowJo (v10.10.0).

### 2.9 Statistical methods

All data are shown as mean ± SEM. Statistical analysis was performed using GraphPad Prism (v10.2.3). Significance was determined by one-way repeated-measures analysis of variance (ANOVA) with corrections for multiple comparisons and subsequent post hoc analysis, as indicated in figure legends. Significance in sex differences in immune proportions (Table 1) were determined with a 2-way ANOVA with Šídák’s post hoc test. For the longitudinal SNI and sham mice leukocyte counts and the longitudinal von Frey and acetone behavior test experiments, because of missing values, significance was determined using the mixed-effects analysis along with subsequent post hoc analysis. A criterion of α=0.05 was used to judge significance.

**Table 1.** Immune cell proportions in male and female mice following sham or SNI surgery.

|  |  |  | Male |  | Female |  | Significance<br>(M vs F) | Statistics<br>(Adjusted<br>p-value) |
| --- | --- | --- | --- | --- | --- | --- | --- | --- |
|  |  |  | Mean % | S.E.M. | Mean % | S.E.M. |  |  |
| Granulocytes | Naïve |  | 20.83 | 2.79 | 8.90 | 0.62 | *** | <0.001 |
|  | Sham | 1 month | 21.23 | 6.97 | 9.40 | 0.74 | ns | 0.1 |
|  |  | 12 months | 14.10 | 1.78 | 19.68 | 1.51 | ns | 0.83 |
|  |  | 18 months | 10.69 | 1.90 | 24.52 | 2.62 | * | 0.01 |
|  | SNI | 1 month | 13.22 | 1.16 | 10.22 | 0.80 | ns | >0.99 |
|  |  | 12 months | 12.90 | 1.28 | 16.98 | 1.02 | ns | 0.96 |
|  |  | 18 months | 11.53 | 1.05 | 18.39 | 1.57 | ns | 0.59 |
| Lymphocytes | Naïve |  | 63.58 | 3.45 | 77.82 | 1.19 | *** | <0.001 |
|  | Sham | 1 month | 65.58 | 7.49 | 78.36 | 1.24 | ns | 0.3 |
|  |  | 12 months | 72.30 | 2.19 | 49.58 | 5.11 | ** | 0.003 |
|  |  | 18 months | 69.43 | 2.60 | 39.77 | 3.54 | *** | <0.001 |
|  | SNI | 1 month | 75.44 | 1.32 | 77.02 | 0.26 | ns | >0.99 |
|  |  | 12 months | 73.82 | 1.96 | 59.80 | 2.61 | ns | 0.16 |
|  |  | 18 months | 73.28 | 2.33 | 50.31 | 5.06 | ** | 0.001 |
| Myeloid | Naïve |  | 13.49 | 0.96 | 10.98 | 1.03 | *** | <0.001 |
|  | Sham | 1 month | 10.67 | 1.59 | 9.32 | 0.71 | ns | 0.3 |
|  |  | 12 months | 11.56 | 0.78 | 27.84 | 4.57 | ** | 0.003 |
|  |  | 18 months | 17.73 | 1.62 | 33.93 | 2.60 | *** | <0.001 |
|  | SNI | 1 month | 8.67 | 0.86 | 9.93 | 0.65 | ns | >0.99 |
|  |  | 12 months | 11.07 | 1.29 | 20.90 | 1.89 | ns | 0.16 |
|  |  | 18 months | 13.63 | 1.82 | 30.26 | 4.59 | ** | 0.001 |

Significance determined by ordinary 2-way ANOVA with Šidák's post hoc test.

## 3. Results

### 3.1 Circulating immune cells display changes related mostly with age but minimally with injury

It is well known that an injury triggers an influx of immune cells to the site of damage to generate local inflammation and promote recovery. However, it has not been fully established whether a local nerve injury would have a long-term impact on circulating immune cells. In this study, we performed sham and sciatic nerve injury (SNI) in young adult (2–3-month old) male and female mice and monitored the number of leukocytes and their sub-groups from 1–18 months post-injury using flow cytometry assay. The gating strategies for each specific cell type are illustrated in Spplementary **Fig. S1**. The absolute cell number was quantified using counting beads. Although no significant differences were detected in the total numbers of circulating leukocytes between SNI and sham mice (**Fig. 1A–B**), we observed age-related regulation in a sexually dimorphic manner. The total number of leukocytes significantly increased in male mice at 18 months following SNI or sham (20 months old), but significantly decreased in female mice starting from at least 7 months post SNI or sham (10 months old).

**Figure 1.**
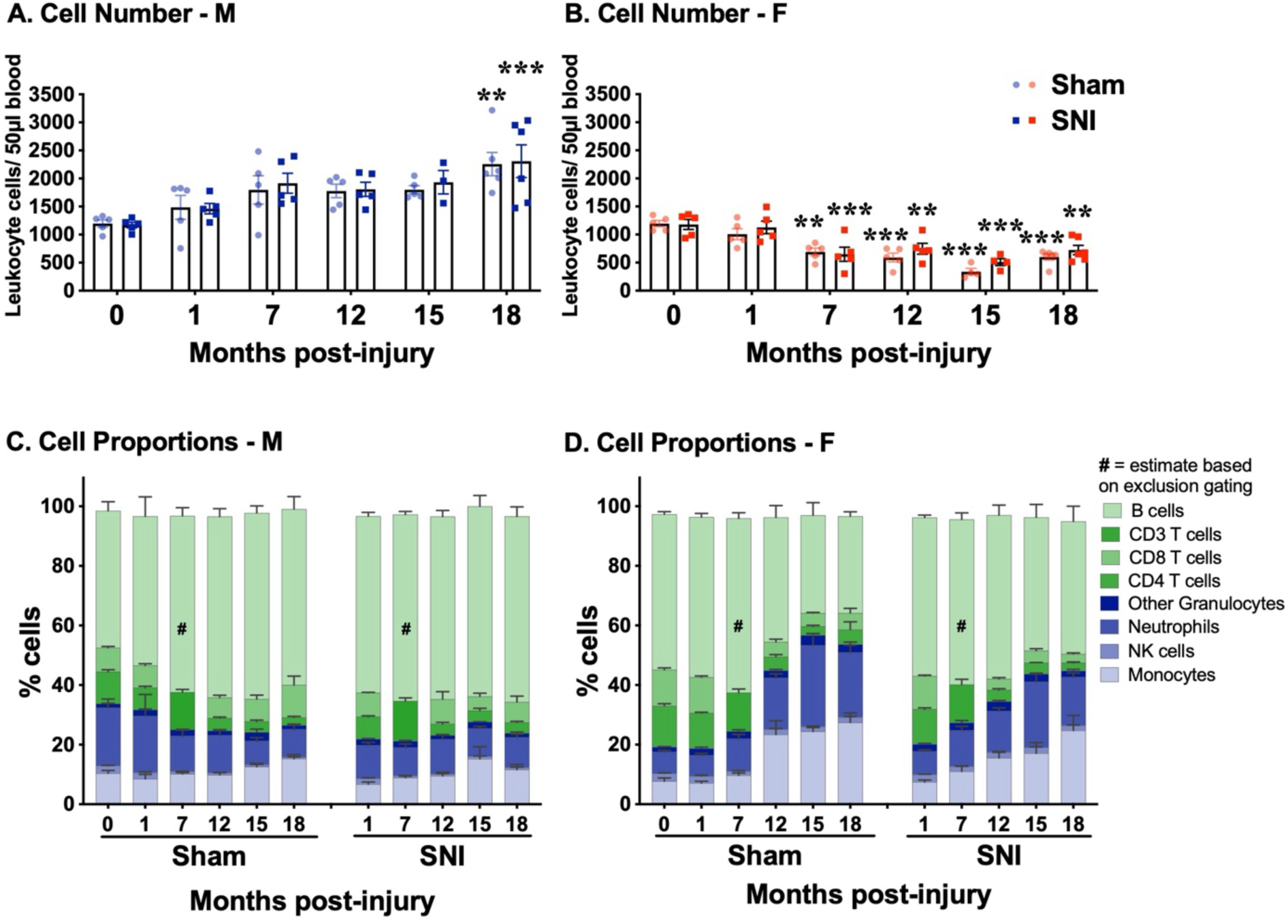
Circulating immune cells display a sex-dependent regulation by age but not by nerve injury. Immune cells were profiled with flow cytometry for 18 months following SNI or sham surgery. Total leukocyte number (50µl blood sample) was quantified with flow cytometry, according to FSC and SSC scatter, for male, sham and SNI **(A)** and female, sham and SNI **(B)**, mice (*n* = 5-6 mice/group). Proportions of each immune cell subset over the total leukocytes were assessed in male, sham and SNI **(C)** and female, sham and SNI **(D)** mice (*n* = 5-6 mice/group). Additionally, blood from several 3-4 months old male and female naïve mice were assayed along with experimental mice at each time point as an internal control, for a total of *n* = 21 male and *n* = 17 female mice. Markers for gating immune cell types are: Monocytes (CD11b, CD115), Neutrophils (CD11b, Ly6G), Other Granulocytes (CD11b, SSC-hi), NK cells (CD11b, NK1.1), CD4 T cells (CD4), CD8 T cells (CD8), and B cells (CD19). **(A-B)** Significance between each time-point and baseline was determined by mixed-effects analysis with Tukey’s post hoc test, \**P*<0.05, \*\**P*<0.01, \*\*\**P*<0.001. All data are presented as the mean ± *SEM*. # indicates that at 7 months post-sham/SNI, CD19 (B lymphocytes) were not included in the panel and the proportion of B lymphocytes were estimated based on exclusion from CD3 and CD11b. Additionally at this time point, CD3, but not CD4 and CD8, was used for assessing T lymphocytes.

In addition to the differential trends in total leukocyte numbers, we also observed a sex difference in immune cell composition in the blood. We first confirmed that before surgery (day 0), female mice had significantly higher proportions of lymphocytes and lower proportions of graulocytes and myeloid cells compared to male mice (**Table 1**). SNI *per se* did not change the immune cell proportions in the blood, either in male or in female mice in any of the examined time points. The proportions of different types of immune cells in male mice maintained relatively stable proportions over the course of aging (**Fig. 1C**). However, as female mice aged, the lymphoid compartment, consisting of B cells, CD4 T cells, and CD8 T cells, contracted progressively over time. The myeloid (mainly monocytes) and granulocyte compartments (mainly neutrophils) expanded proportionally (**Fig. 1D**). Both lymphocytes and myeloid cells were significantly different between male and female mice at 18 months following either sham or SNI surgery (**Table 1**). Overall, changes observed in immune cell numbers and proportions appear to be mostly related to age and are minimally impacted by SNI. Statistical difference of granulocyte, myeloid, and lymphoid cells in male and female mice across the time points of 1, 12, and 18 months is listed in **Table 1**.

### 3.2 Serum mediators in mice are regulated by nerve injury in a sexually dimorphic manner

As we did not detect injury-specific changes in circulating immune cells, we then investigated the impact of a local nerve injury on molecular mediators in the serum. By using the Proteome Profiler Assay, we assessed 111 inflammation-related mediators and growth factors in the serum of young naïve male (**Fig. 2A**) and female (**Fig. 2B**) mice. Data were analyzed and plotted as Log_2_ of fold changes (male vs. female). As depicted in **Fig. 2C**, among all examined targets, around one-third of the molecules were higher in male, while two-thirds of the molecules were higher in female. The quantified data shows that the top three proteins more abundant in naïve male serum are the chemokines CCL-11, CCL-21, and CXCL-13, whereas the top three proteins more abundant in naïve female serum are the soluble vascular cell adhesion molecule-1 (VCAM-1), the complement component C5/C5a, and preadipocyte factor-1 (Pref-1), although it is worth noting that the absolute levels of some targets may not be particularly high. The functional categories of the serum proteins are displayed in **Table 2**. Overall, female mice have more serum proteins with higher levels than those in males, in both immune-related and non-immune related-protein categories. It appears that in physiological conditions, males have more proteins related to chemotaxis, whereas female mice have more cytokines, complements and pentraxins. In all, it appears that serum proteins of healthy young mice display sex-dependent features. To ensure the data was not biased by potential outliers in pooled samples, we verified the levels of some targets such as G-CSF, TNF-α, CCL-2, and CCL-4 from individual serum samples of naïve young male and female mice in a Luminex assay, Data was depicted in Supplementary **Fig. S2**.

**Figure 2.**
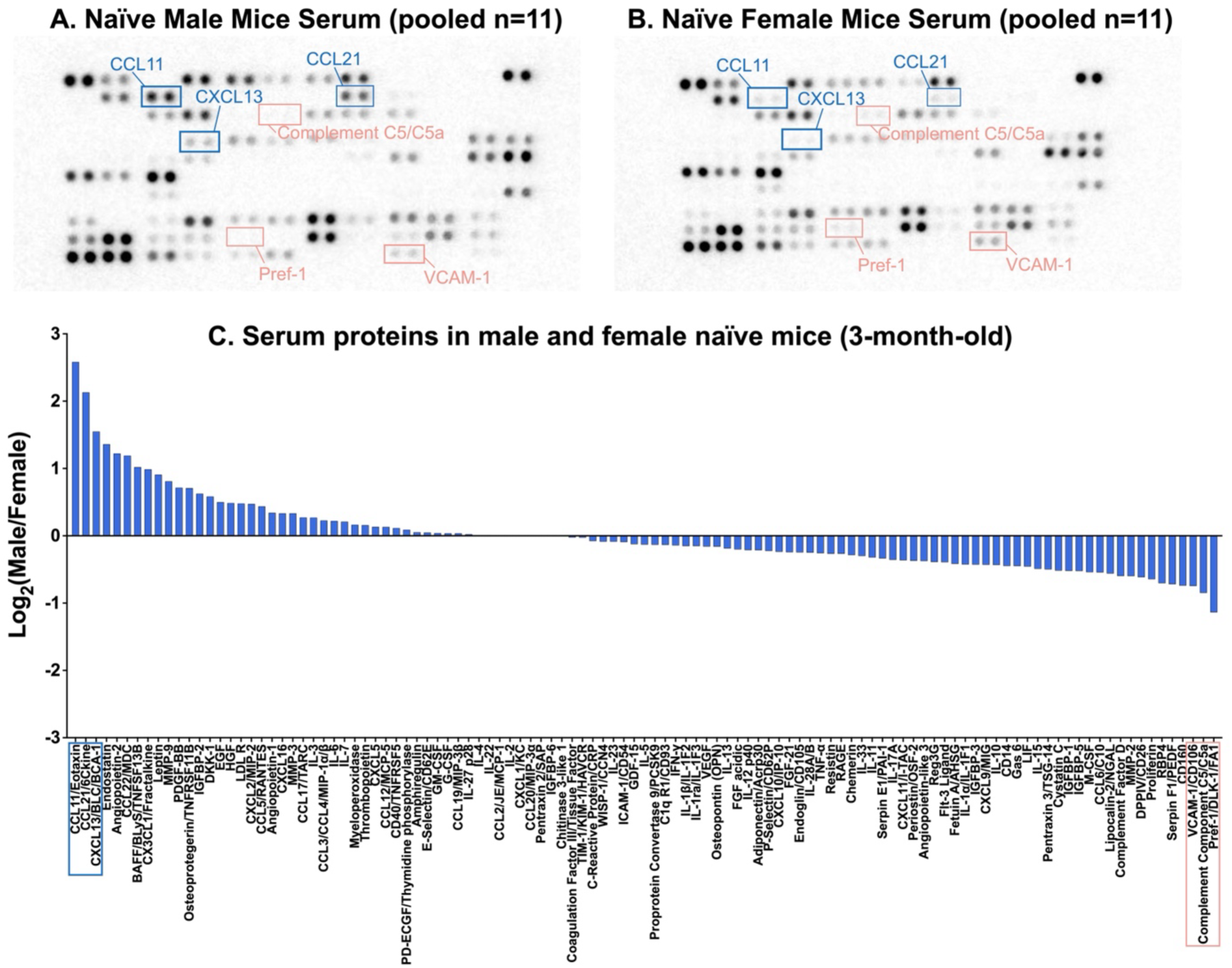
Serum protein composition is different between young naïve male and female mice. Proteome Profiler blots with pooled serum from naïve male **(A)** and female **(B)** mice. Blue boxes indicate the top 3 proteins that are higher in male compared to female mouse serum. Peach boxes indicate the top 3 proteins higher in female compared to male mouse serum. Quantification of the log_2_ fold change of 3 months old naïve male over 3 months old naïve female was plotted for each serum target in Proteome Profiler blots **(C)**. The targets are arranged from highest to lowest concentration in male compared to female mouse serum. For **A-C**, serum samples were pooled (n=11/group).

**Table 2.** Serum proteins in naïve male and female mice are classified into functional categories.

| **Mediator Categories** | **Number of proteins higher in Male young naïve** | **Number of proteins higher in Female young naïve** |
| --- | --- | --- |
| **Immune-related proteins** | **29** | **38** |
| *Pro-inflammatory cytokines* | *5* | *14* |
| *Anti-inflammatory cytokines* | *2* | *7* |
| *Adaptive immune response cytokines* | *5* | *4* |
| *Chemotaxis* | *17* | *7* |
| *Complement factors* | *0* | *3* |
| *Pentraxins* | *0* | *3* |
| **Non-immune-related proteins** | **16** | **28** |
| *Growth factors* | *8* | *10* |
| *Coagulation factors* | *0* | *2* |
| *Metabolic factors* | *2* | *9* |
| *ECM mediators* | *3* | *2* |
| *Bone remodeling* | *2* | *2* |
| *Angiogenesis factors* | *1* | *3* |

By using the same Proteome Profiler Assay containing 111 targets, we next investigated changes in serum proteins at 1, 12, and 20 months post-surgery (corresponding to 4, 15, and 23 months-old) in male and female, sham and SNI mice. Data were first analyzed as Log2 fold changes (sham vs. 3 months old naïve, and SNI vs. 3 months old naïve) within their respective sex- and age-matched groups. In male mice, at one month post-sham surgery, more than half of the inflammation-related proteins were upregulated (**Fig. 3A**). This upregulation was further amplified by age, which appears to have peaked at 12 months post-surgery when the mice were around 14-months old. At one month post-SNI, serum from male SNI mice have a larger number of inflammatory proteins with increased levels compared to that of the sham group. It is obvious that SNI induced systemic inflammation in male mice, and such disturbances were maintained across the lifespan (up to 20 months post-injury, 22-months old). Very surprisingly, we did not detect similar changes in female mice, either in the sham or SNI groups (**Fig. 3B**). Sham surgery resulted in an upregulation of most serum proteins in female mice, which were different from those in male mice, and was maintained at relatively stable levels over 2 years. SNI did not exacerbate systemic inflammation, but rather generally reduced it in females, opposite to what was observed in male mice. To confirm the pattern of injury-induced systemic inflammation, we performed a Luminex assay on some circulating molecular targets with individual samples of separate serum sets, which were from 6 and 15 months post-sham and SNI surgery male and female mice, where we observed similar sexual dependent regulation. Compared with naïve mice, sham surgery/SNI triggered a 1-3 Log2 fold increase of CCL-4, CCL-2 and TNF-α in male, but not in female mice at two examined time-points, although the difference between SNI and sham injured mice was not detectable in these specific inflammatory targets (**Fig. 3C**), which may be due to the interfluences by age-associated systemic inflammation. The direct impact of SNI on serum immune-related protein levels was further confirmed by subtracting the sham surgery-induced levels of the same proteins in the Proteome Profiler data. As depicted in **Fig. 4**, SNI induced an increase of most examined systemic inflammatory proteins across the lifespan in male mice (**Fig. 4A**), but inhibited the release of inflammatory molecules in the serum of female mice compared to that of sham (**Fig. 4B**).

**Figure 3.**
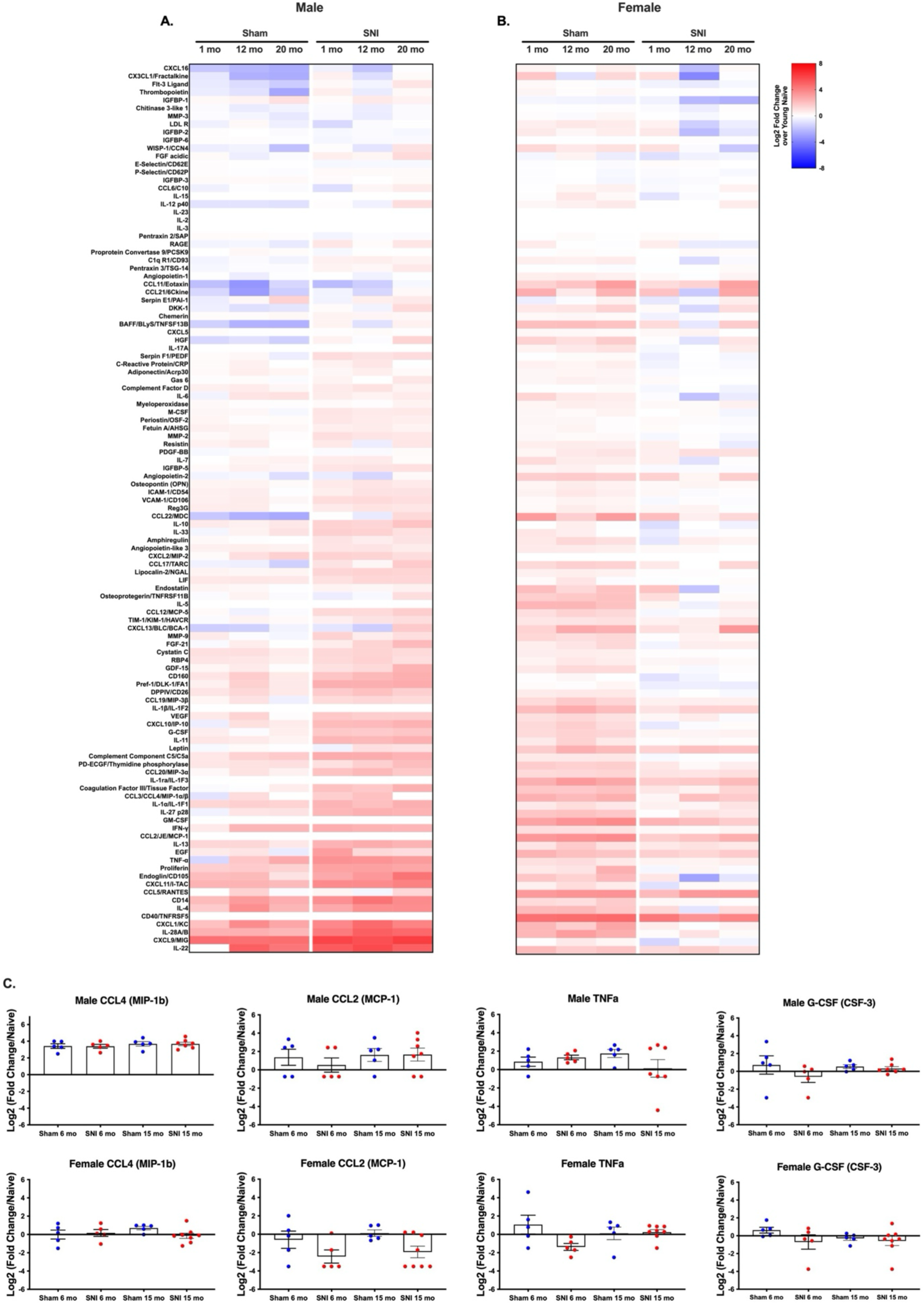
Sham- and SNI-surgeries have long-term impact on serum protein composition in male and female mice. The release of inflammation-related serum proteins is increased in sham-operated mice, while nerve injury (SNI) accentuates the upregulation of these proteins in male mice but downregulates some in female mice, and such impact persists across the lifespan. Quantification of Proteome Profiler blots of male **(A)** and female **(B)**, sham and SNI mice over 3 time points (1-month, 12-months, and 20-months). The data is plotted as a log_2_ fold change for each condition (sham and SNI, male and female) at each time point (1, 12, 20 months) over the 3 months old sex-matched naïve mouse serum. Protein targets are arranged in order from lowest to highest, sorted based on each serum protein target’s average log_2_ fold change across all conditions. For **A-B**, serum samples were pooled (naïve: *n* = 11 mice/group; 1-month: n = 9 – 10 mice/group; 12-months: n = 7 – 9 mice/group; 20-months: 3 – 9 mice/group). Luminex multiplex immunoassay quantification of individual serum samples taken from 6 and 15 months time points following sham or SNI surgery **(C)**. Data for the Luminex plots are presented as the mean ± *SEM* of a log_2_ fold change for each condition over the average 3 months old sex-matched naïve mouse serum. *n* = 5-8 mice/group.

**Figure 4.**
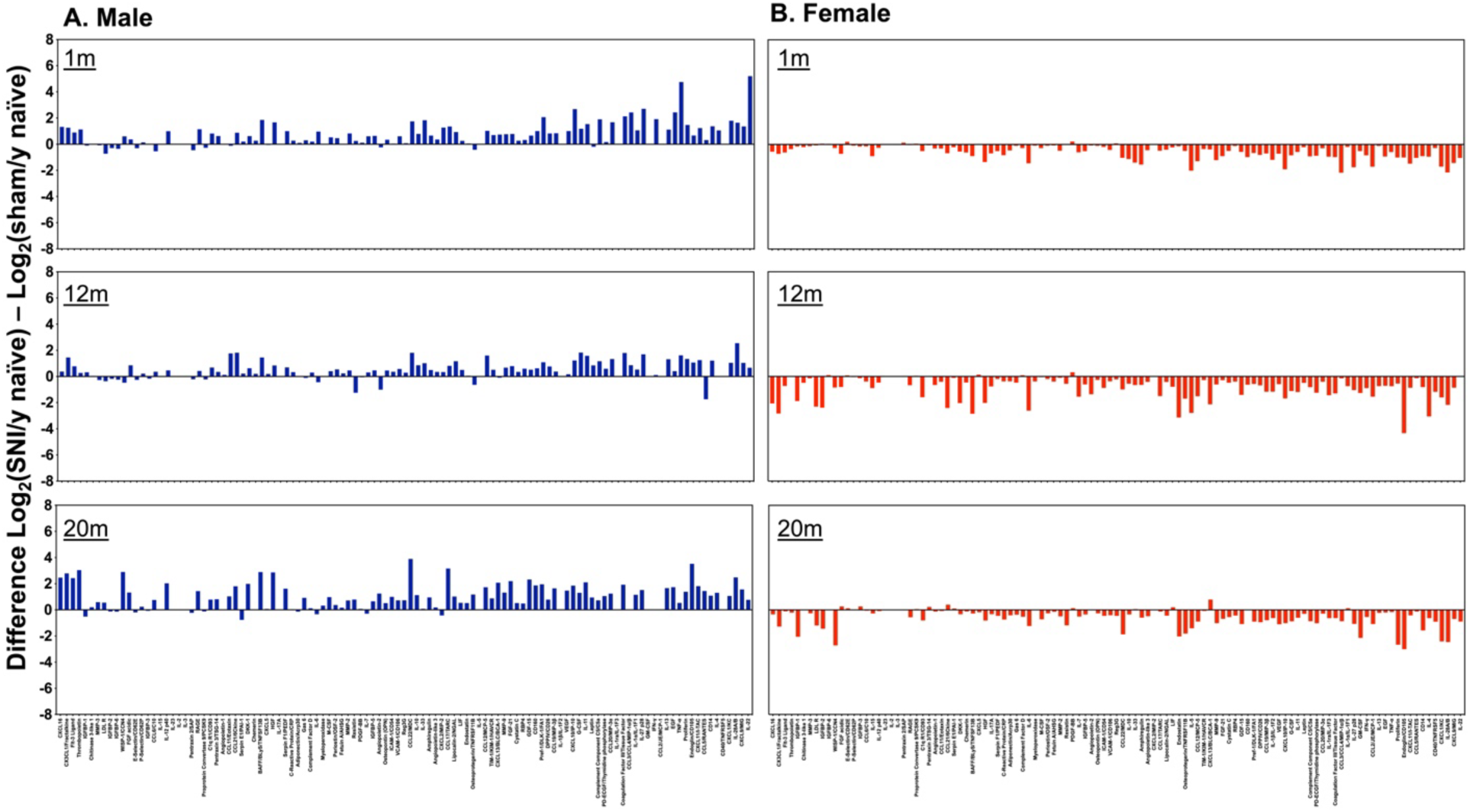
Compared to sham-surgery, SNI up-regulates most serum proteins in male but down-regulates them in female mice. The difference between serum targets in SNI and sham mice was calculated by subtracting the log_2_ fold change of SNI mice over 3 months old sex-matched naïve mice from the log_2_ fold change of sham mice over 3 months old sex-matched naïve mice. The time points at 1-month, 12-months, and 20-months are shown for male **(A)** and female **(B)** mice. The same order of protein targets from Fig. 3 was applied in the graph.

### 3.3 Sexual dimorphism in mouse serum persists regardless of regulation by age or nerve injury

To further elucidate the patterns of inflammation and sexual dimorphism in the sham and SNI mouse serum, we combined the aggregated data from the different time points in both nerve injured and sham mice to identify proteins that are consistently released in either male or female mice. As shown in **Fig. 5A**, in sham and SNI mice, across different ages and injury conditions, several proteins including the soluble glycoprotein CD160 receptor, chemokines (CXCL-9 and CXCL-11), and insulin growth factor binding proteins (IGFBP1 and IGFBP3) are consistently higher in abundance in male mice than in female mice. On the other hand, the top proteins that are consistently higher in abundance in female than in male mice are proteins related to vasculature or lipid metabolism such as angiopoietins (Ang1 and Ang2) and leptin, chemokines (CCL-5, CCL-11, and CXCL-13), as well as factors involved in tissue regeneration like the platelet-derived growth factor-BB (PDGF-BB) (**Fig. 5B**). The serum proteins observed in **Fig. 5A–B** for the analysis of sex differences are grouped into functional categories and shown in **Table 3**. Overall, across different age (4, 15, 23 month-old) and injury conditions (sham or SNI), male mice have higher levels of immune-related serum proteins, whereas female mice have higher non-immune related proteins in the serum (**Table 3**). In the context of aging, we aggregated data from both sham and SNI mice, still separated by sex, to identify serum proteins that steadily increased as the mice aged. The plasminogen activator inhibitor (PAI-1), encoded by *Serpin E1*, is a serine protease inhibitor displaying a consistent, age-dependent increase in both male and female mice (**Fig. 5C–D**). The chemokine CCL-6, however, was observed to steadily increase with age only in female mice (**Fig. 5D**). The serum proteins observed in **Fig. 5C–D** for the analysis of aging sex differences are grouped into functional categories and shown in **Table 4**. When we aggregated data across all ages to analyze the sex differences resulting from nerve injury, we observed a striking difference in the impact of nerve injury on male and female mouse serum homeostasis. In male mice, compared to sham surgery across all ages, SNI induced the upregulation of 55 serum proteins, most of which are inflammation-related cytokines and chemokines (**Fig. 5E**, **Table 5**). However, in female mice, only 3 serum proteins, including one chemotaxis-related molecule, the soluble adhesion molecule E-selectin, was consistently upregulated in SNI mice compared to sham mice regardless of age (**Fig. 5F**). The serum proteins observed in **Fig. 5E–F** for the analysis of sex differences due to nerve injury are grouped into functional categories and shown in **Table 5**. This analysis further confirmed that the impact of nerve injury on the immune system across the lifespan is sexually dimorphic.

**Figure 5.**
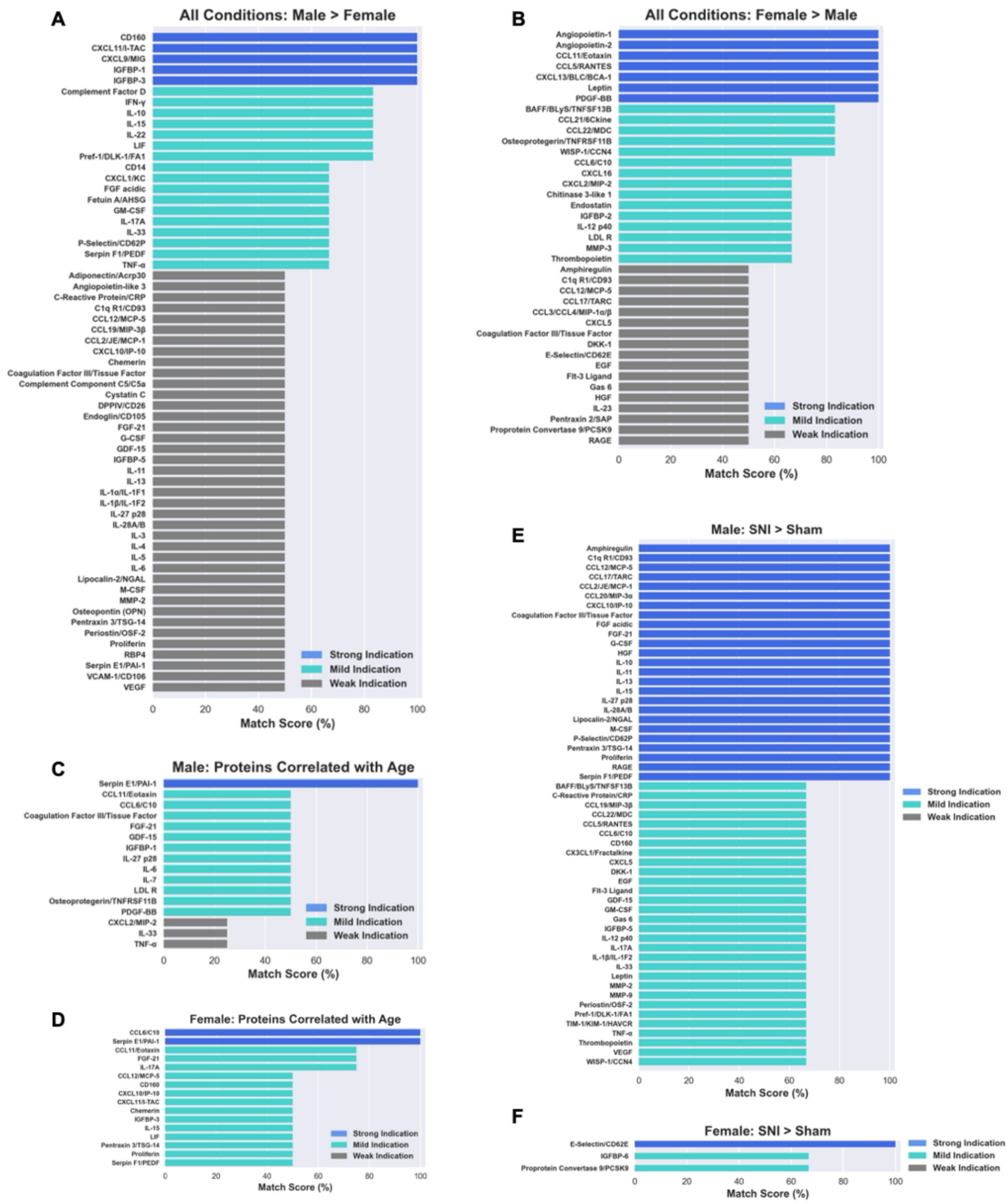
Sexual dimorphism in mouse serum persists regardless of regulation by age or nerve injury. All conditions (nerve injury status, age) were aggregated to identify mediators consistently higher in male **(A)** or female **(B)** mice. Sham and SNI data were aggregated to identify mediators that increased across age in male **(C)** and female **(D**) mice. Data across all ages were aggregated to identify mediators that were increased by SNI compared to sham in male **(E)** and female **(F)** mice.

**Table 3.** Sex difference of top proteins in the serum of male and female mice regardless of regulation by age or nerve injury (analyzed from Fig. 5A**-B**).

| **Sex Difference – strong to mild indication only** |  |  |
| --- | --- | --- |
| **Mediator Categories** | **M>F** | **F>M** |
| **Immune-related proteins** | **16** | **11** |
| *Pro-inflammatory cytokines* | *6* | *3* |
| *Anti-inflammatory cytokines* | *3* | *0* |
| *Adaptive immune response cytokines* | *2* | *0* |
| *Chemotaxis* | *4* | *8* |
| *Complement factors* | *1* | *0* |
| *Pentraxins* | *0* | *0* |
| **Non-immune-related proteins** | **6** | **11** |
| *Growth Factors* | *3* | *3* |
| *Coagulation factors* | *0* | *0* |
| *Metabolic factors* | *3* | *2* |
| *ECM mediators* | *0* | *2* |
| *Bone remodeling* | *0* | *2* |
| *Angiogenesis factors* | *0* | *2* |

**Table 4.**
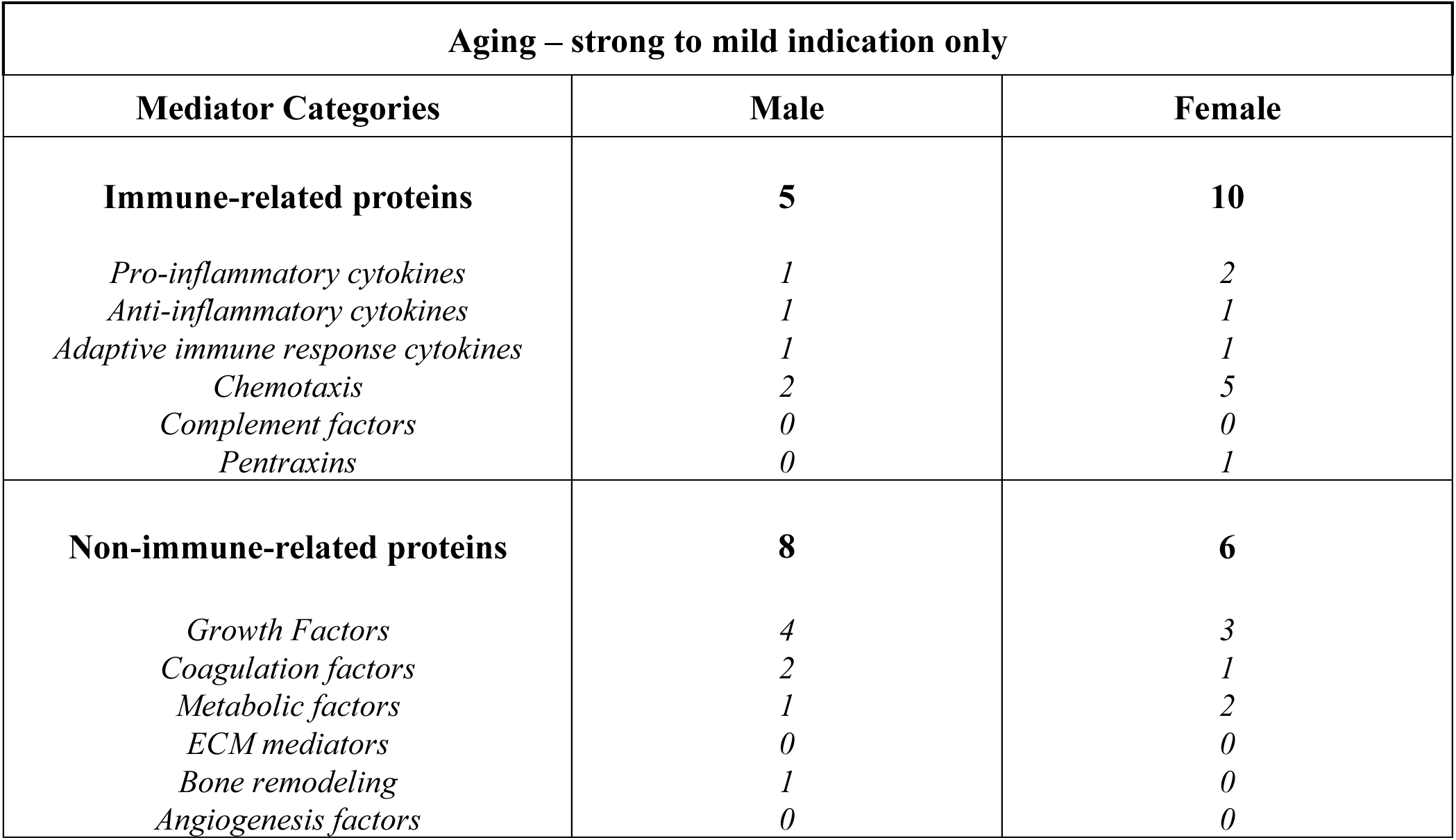
Top proteins from analysis of aggregate data for aging-related targets (Fig. 5C**-D**) are classified into functional categories.

| **Aging – strong to mild indication only** |  |  |
| --- | --- | --- |
| **Mediator Categories** | **Male** | **Female** |
| **Immune-related proteins** | **5** | **10** |
| *Pro-inflammatory cytokines* | *1* | *2* |
| *Anti-inflammatory cytokines* | *1* | *1* |
| *Adaptive immune response cytokines* | *1* | *1* |
| *Chemotaxis* | *2* | *5* |
| *Complement factors* | *0* | *0* |
| *Pentraxins* | *0* | *1* |
| **Non-immune-related proteins** | **8** | **6** |
| *Growth Factors* | *4* | *3* |
| *Coagulation factors* | *2* | *1* |
| *Metabolic factors* | *1* | *2* |
| *ECM mediators* | *0* | *0* |
| *Bone remodeling* | *1* | *0* |
| *Angiogenesis factors* | *0* | *0* |

**Table 5.**
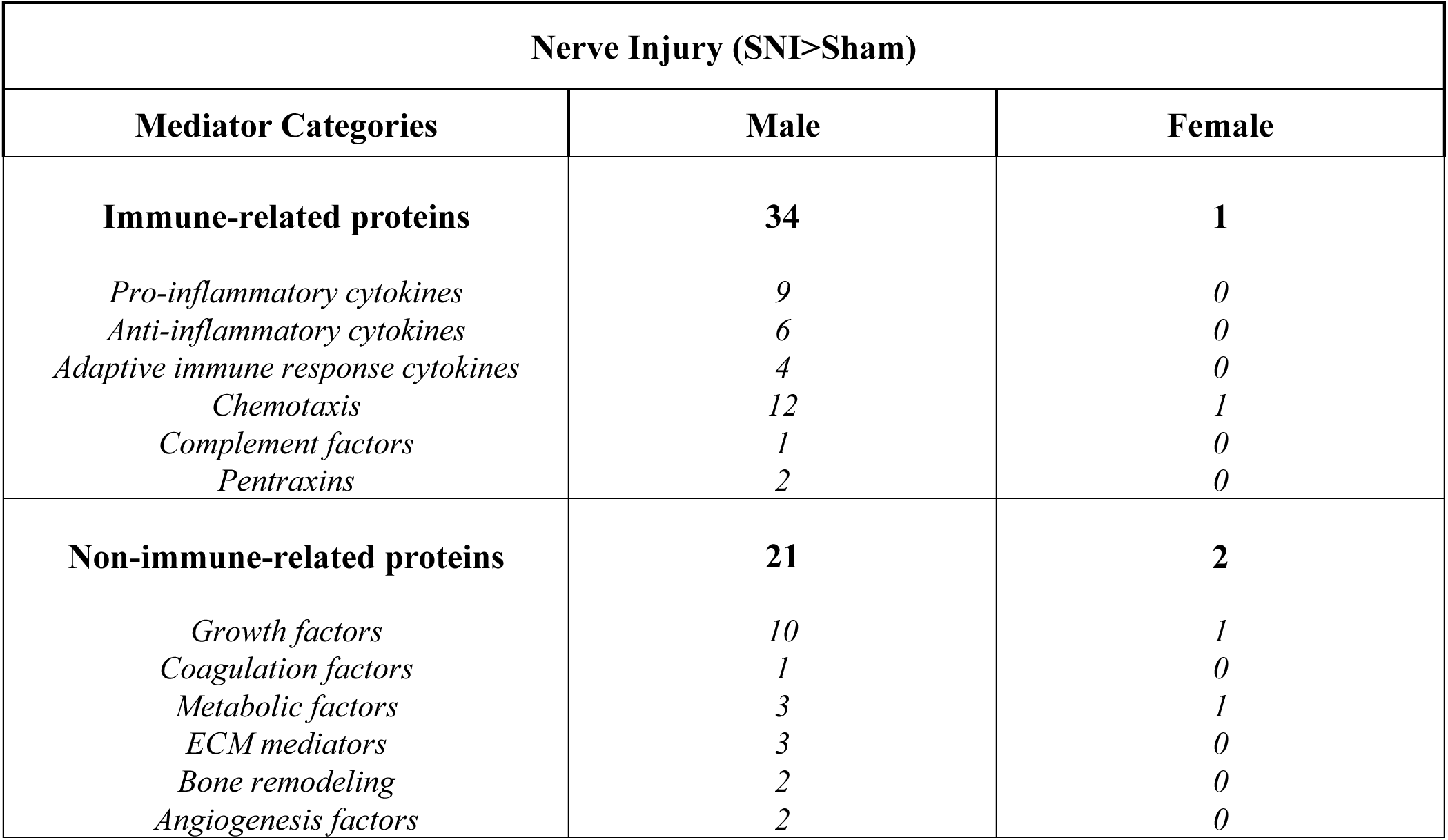
Top proteins from analysis of aggregate data for nerve-injury-related targets (Fig. 5E**-F**) are classified into functional categories.

### 3.4 Nerve injury induced systemic disturbance triggers painful response in both male and female naïve mice

Nerve injury often (although does not always) result in neuropathic pain. We confirmed that in conjunction with the systemic immune changes, SNI induced-neuropathic pain lasted for at least two years in both male and female mice. Male mice developed stable and long-lasting mechanical allodynia (**Fig. 6A**) and progressively increasing cold hypersensitivity (**Fig. 6B**). Similar patterns were observed in female mice (**Fig. 6C-D**).

**Figure 6.**
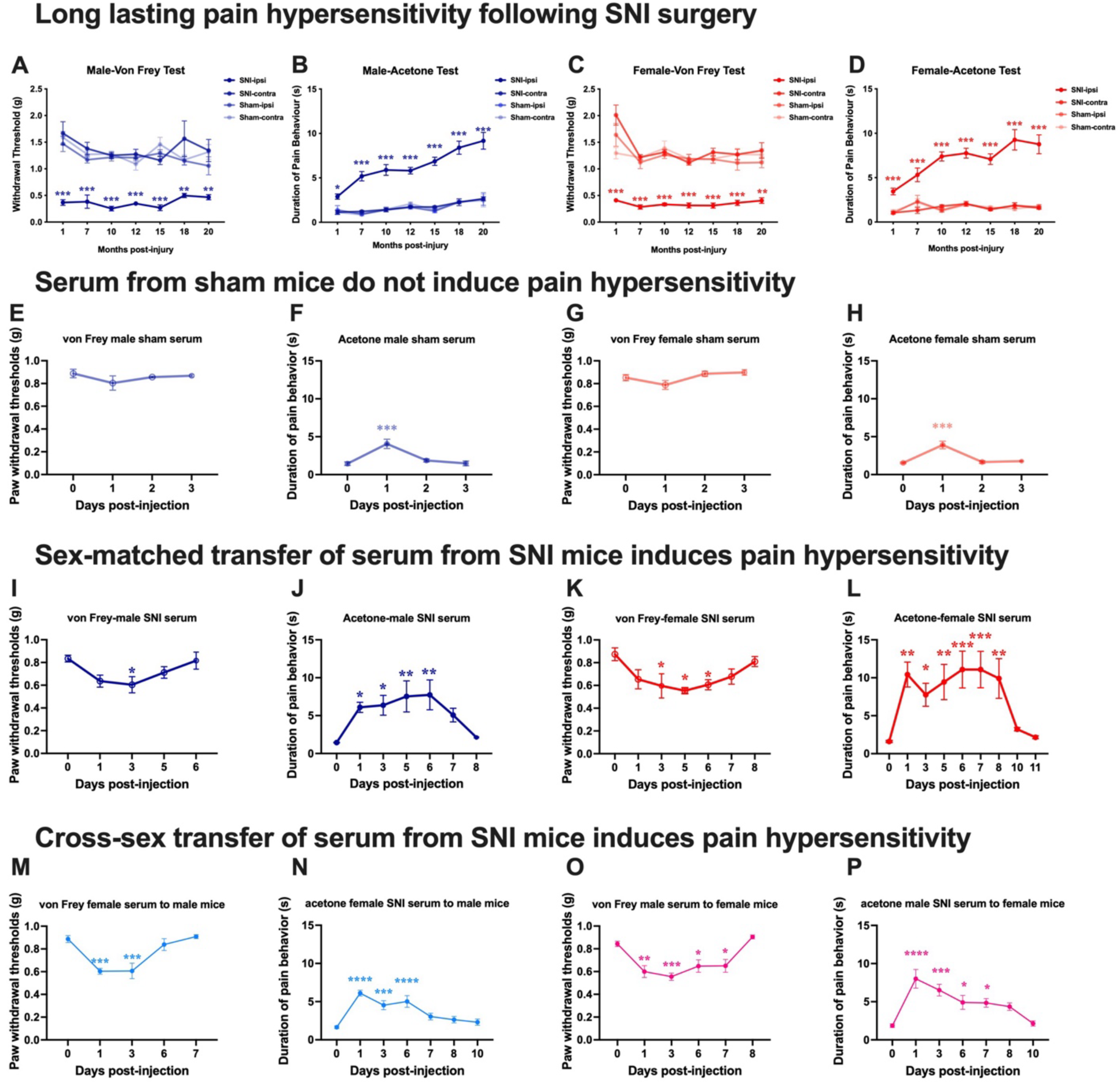
Serum from both male and female nerve injured mice induces mechanical and cold hypersensitivity in young naïve mice. **A-D:** Long lasting pain hypersensitivity following SNI surgery. Male mice von Frey **(A)** and acetone **(B)** tests and female mice von Frey **(C)** and acetone **(D)** tests over 20 months following sham or SNI surgery (*n* = 3-10 mice/group/time point). **E-H:** Serum from sham mice at one month post-surgery do not induce pain hypersensitivity. Male recipient mice von Frey **(E)** and acetone **(F)** tests, and female recipient mice von Frey **(G)** and acetone **(H)** tests following sham serum injection (100µL/injection; *n* = 5 mice/group). **I-L:** Sex-matched transfer of serum from SNI mice at one month post-surgery induces pain hypersensitivity in both male and female young naïve mice. Male mice von Frey **(I)** and acetone **(J)** tests, and female mice von Frey **(K)** and acetone **(L)** tests following SNI serum injection (100µL/injection; *n* = 5 mice/group). **M-P:** Cross-sex transfer of serum from SNI mice at one month post-surgery induces pain hypersensitivity in both male and female young naïve mice. von Frey **(M)** and acetone **(N)** tests in male recipient mice, and von Frey **(O)** and acetone **(P)** tests in female recipient mice. (100µL/injection; *n* = 5 mice/group). All tests were performed until pain sensitivity returned to pre-serum transfer levels. Significance between the behavior measurement in the SNI ipsilateral paw and in the sham ipsilateral paw was assessed by mixed-effects analysis with Tukey’s post hoc test in **A-D**. Significance between the behavior measurement at each time point and at baseline was assessed by mixed-effects analysis with Dunnett’s post hoc test in **I**, and one-way repeated-measures ANOVA with Dunnett’s post hoc test in **E-H** and **J-P**, \**P*<0.05, \*\**P*<0.01, \*\*\**P*<0.001, ****P<0.0001. All data are presented as the mean ± *SEM*.

To understand the functional consequence of the SNI-associated, long-lasting systemic disturbance, we intravenously transferred serum from 1 month post-SNI or -sham mice to naïve mice to determine the potential of systemic contribution to pain. While sham serum from male mice did not induce any mechanical allodynia (**Fig. 6E**), it induced a transient and slight cold hypersensitivity lasting for 1 day (**Fig. 6F**) when transferred to male naïve mice. The exact same pattern was detected in female mice (**Fig. 6G-H**). On the other hand, when serum from male 1 month post-SNI mice was transferred to sex-matched naïve male mice, the recipient mice developed mechanical allodynia peaked at day 3 of post-injection (**Fig. 6I**) and cold hypersensitivity lasted for 6 days (**Fig. 6J**). Surprisingly, when serum from female 1 month post-SNI mice was transferred to sex-matched naïve male mice, it also induced 6 days of mechanical allodynia (**Fig. 6K**) and 8 days of cold hypersensitivity (**Fig. 6L**). Statistical difference of sham- or SNI-serum induced changes in mechanical and cold sensitivity of naïve mice (day 1 – day 3) is illustrated with area under the curve analysis in Supplementary **Fig. S3**. Interestingly, cross-sex administration of 1 month post-SNI serum also induced pain sensitivity in naïve mice. Serum from female SNI mice induced mechanical allodynia (**Fig. 6M**) and cold hypersensitivity (**Fig. 6N**) in male naïve recipient mice. Vice versa, serum from male SNI mice triggered both mechanical (**Fig. 6O**) and cold allodynia (**Fig. 6P**) when administered to female naïve mice. SNI serum (transferred through intravenous administration)-triggered hypersensitivity is more likely generalized pain as we detected on both paw, which was similar to our previous report that serum from nerve injured mice induced mechanical hypersensitivity occurring not only in the paw, but also in the orofacial region (10). These results demonstrate that SNI disturbed serum homeostasis in both male and female mice. Regulated serum proteome could contribute to enhance pain sensitivity. Even though only male SNI mice serum had robust increases in various serum cytokines and chemokines, female SNI mice serum still triggered the development of pain hypersensitivity in not only sex-matched female naïve mice, but also in male naïve mice.

## 4. Discussion

Nerve injury can trigger inflammation both at the site of the damage and remotely in the DRG and the spinal cords (4, 22). However, it has not been fully established whether such an insult could result in systemic disturbances. In an attempt to fill this knowledge gap, in this study we assessed the molecular and cellular changes in the blood of both male and female mice following a painful nerve injury across the lifespan. We demonstrated that spared sciatic nerve injury-associated systemic disturbance persists, along with neuropathic pain, over two years. Our data also revealed that the immune system of male and female mice respond differently to the nerve injury. Among 111 tested molecular targets, nerve injury-induced upregulation occurred in male mice but not in female mice. Surprisingly, regardless of what changes were detected in the serum, both male and female serum from nerve-injured mice were capable of inducing pain hypersensitivity when transferred to naïve mice, suggesting distinct mechanisms for the serum to elicit pain in female and in male mice.

### 4.1 Sex difference in the immune system of naïve mice

Sexual dimorphism in the immune response has been extensively studied; women usually exhibit lower infection rates than men but have higher incidence of autoimmune diseases compared to men (19, 23). However, the baseline profile of serum protein compositions, especially cross-sex comparisons, has been barely reported. The sex-specific pattern of serum protein distribution observed in our mouse study is indeed in line with previous human reports, where proteins involved in immune cell chemotaxis are more abundant in men, and other molecules involved in non-immune related function, such as fatty acid oxidation/hormone regulation, cell growth/death have been found at higher levels in women (24). We do recognize that our classification of serum proteins into specific functional categories may be oversimplified, given that many of these molecules are pleiotropic and participate in multiple inflammatory pathways, depending on context and the nature of the immunological insult (25). Some proteins in our non-immune-related category may be indirectly involved in immune regulation (24). Similar to previous reports in humans (26, 27) and in mice (28), we also confirmed the sex differences in circulating immune cells, specifically higher proportions of both B and T lymphocytes, and lower proportions of myeloid cells in female compared to male mice.

### 4.2 Sex differences in age-related changes in serum proteins

In patients, neuropathic pain can sometimes persist for the rest of their life, even with treatment. To investigate more specifically the long-term consequences of nerve injury on the immune system, we monitored mice for 20 months post-SNI surgery. Thus, inevitably, our data not only depict changes related to injury, but also related to aging. During the normal aging process, the activity and function of the immune system declines, known as immune senescence. At the same time, aging individuals also develop inflammaging: a chronic, systemic, low-grade, and subclinical proinflammatory state (29). Inflammaging is represented by an enhanced secretory profile, the senescence-associated secretory phenotype (SASP), through which senescent cells modulate their local environment (30). Although we all age with inflammatory signatures, data on sex differences in inflammaging, especially a comparison of serum proteome features in male and female aged individuals is missing. In addition to confirming that only female mice have contraction of T/B lymphocytes and expansion of myeloid/granulocytes with age, in this study we also identified age-specific changes in serum mediators that feature sexual dimorphism. While Serpin E1/PAI-1, a key cellular senescence marker (31), is consistently associated with aging in both male and female mice across all conditions, the increase of CCL-6/C10, a potentially neuroprotective molecule (32), was only consistent in female mice. It is worth emphasizing that this set of data was not generated with naïve mice. Instead, all mice were challenged by either sham surgery or SNI at the young-adult stage of their lives, which could produce factors contributing to the development of inflammaging. In the case where no apparent difference was found between SNI and sham mice, the upregulation of molecules in the serum vs their young sex-matched counterparts could be considered as a sign of aging-associated systemic inflammation rather than a phenomenon specifically induced by nerve injury.

### 4.3 Sex differences in nerve injury-related serum disturbances

Most of our knowledge on whether peripheral nerve injury can lead to systemic inflammation comes from human studies. Among them, many are in the field of pain research where the increase of several inflammatory cytokines has been linked to nerve injury-associated neuropathic pain in patients (7, 8, 33). Unfortunately, none of these studies have separately analyzed male and female patient data. Furthermore, very few animal studies have investigated systemic alterations following nerve injury. Most of them (11, 34, 35), including our previous report (10), have worked with male rats/mice. Only one mouse study used both male and female animals and reported sex difference in immune cell trafficking, and on the levels of inflammatory mediators in the blood and in the injured nerve (36). Information on sex differences in injury-induced changes in the serum/plasma is lacking. To our knowledge, the current study is the first to assess both longitudinal changes and sex differences in serum factors resulting from nerve injury. Our observations are that in male mice, following SNI, a systemic inflammation developed. Across all conditions (nerve injury status, age), a significant number of serum mediators with 34 immune-related proteins and 21 non-immune related proteins were upregulated. Surprisingly, only one immune-related protein and two non-immune-related proteins are upregulated in female nerve-injured mice. Another interesting phenomenon that has drawn our attention is that even sham surgery could lead to the regulation of some serum proteins, which is again long-lasting. Such phenomenon was specifically obvious in female mice. Although sham surgery-induced systemic alteration occurred in both male and female mice, the regulated serum proteins were not the same. Taken together, the data highlight the importance of injury—whether it is minor, like sham surgery, or severe, like SNI—in systemic homeostasis. The impact could be long-lasting, even across the lifespan, yet with a distinctive pattern in male and female mice.

### 4.4 Functional significance of nerve injury/sham surgery triggered systemic disturbance

Although more and more clinical evidence indicates that there may be an association of increased levels of several cytokines/chemokines in the blood with nerve injury/trauma-triggered painful neuropathy (37), the causal relationships still need to be confirmed. Our serum transfer experiments attempted to fill this gap. It is interesting to note that although serum factors in both sham and SNI mice are regulated compared to naïve mice, only the SNI (one-month post injury) serum induced both mechanical allodynia and cold hypersensitivity in naïve recipients, whether transferred in a same-sex or cross-sex manner. The findings suggest that although a minor injury like a sham surgery also generates systemic chronic inflammation, the composition and the intensity of such reactions are different from that triggered by a direct injury to peripheral nerve. Thus, when mice are young, the systemic changes after minor injury may not have significant, or have very mild, transient functional consequences, at least in terms of sensation, whereas SNI-triggered systemic disturbances can alter mouse sensory functions. Serum from SNI mice elicited pain hypersensitivity in naïve mice, suggesting that SNI-associated changes in serum could be a contributor to neuropathic pain. However, as mice age, the situation may change. We have observed that serum from normal aging mice can decrease pain thresholds in young mice recipeints (data not shown). It is likely that serum from aging sham mice also contributes to modifying pain sensitivity, potentially due to the development of inflammaging as the mice become older. Additionally, the upregulation of inflammatory mediators in the systemic environment of aging sham and SNI mice suggests a potential overlap between inflammaging, which is a part of the normal aging process, and chronic systemic inflammation resulting from nerve injury. This prolonged chronic inflammation may have far-reaching consequences even beyond the context of pain, serving as risk factors for a range of chronic non-communicable diseases including cardiovascular diseases, autoimmunity, and neurodegenerative disorders (38).

The involvement of cytokines/chemokines in neuropathic pain has been extensively investigated in the past and their roles are well established (39–41). In the serum of male SNI mice, an increase of a large group of immune-related proteins including cytokines and chemokines were consistently detected in our study. These serum proteins could be important players in eliciting hypersensitivity when transferred to naïve mice. They should also be contributors to the development and maintenance of neuropathic pain in nerve-injured male mice. However, additional investigations are needed to determine the contribution of cytokines and chemokines, especially these proteins have short half-life but SNI-serum induced hypersensitivity last few days. The potential of autoantibodies could be considered as they have much longer half-life, although we didn’t detect significant changes in the overall levels of any immunoglobulin classes in our SNI serum samples (data not shown). Besides, the possibility of secondary immune activation following acute cytokine/chemokine stimulation cannot be excluded. Similarly, the key culprits in the serum of female nerve-injured mice responsible for inducing pain hypersensitivity in naïve mice remains to be determined. The Proteome Profiler Mouse XL Cytokine Array Kit (ARY028, R&D systems) that was used in this study allowed us to simultaneously detect 111 soluble mouse proteins, including cytokines, chemokines, and growth factors. However, there are many other proteins in the serum, such as albumin, antibodies, and many non-protein components, such as lipids, hormones, metabolites, extracellular vesicles, etc. Additional investigations are needed to identify pain hypersensitivity-inducing factors in the serum of female nerve-injured mice, which may not be exclusively proteins but possibly even metabolic components such as lipids or carbohydrate molecules or moieties with signaling properties.

### 4.5 Limitations of the study

There are several limitations in the current study. First, by pooling the serum samples in this study, we were only able to measure the central tendency of each group of mice and were unable to assess the spread of the data. However, we were able to reproduce similar data showing a sex difference in serum proteins at two time-points using individual samples in a Luminex assay. Second, while Proteome Profiler Mouse XL Cytokine Array Kit allowed us to examine the involvement of more than 100 serum proteins across age and different injury conditions, it is not an unbiased approach. Thus, we might miss other targets that are not included in the kit. Third, although we revealed a sex-dependent pattern of SNI triggered systemic inflammation, it is difficult for us to identify and validate specific factor(s) responsible for hypersnesitivity. In these complicated biological processes, multiple cytokines and chemokines operate in a network, such that the action of each single cytokine or chemokine could be regulated by the presence or absence of others. Especially in our case where the systemic inflammation we are dealing with is a sterile, low-grade inflammation, we don’t believe that any one specific target(s) will be acting alone at a significantly higher level compared to other molecules; rather it is likely multiple molecules/factors acting together to synergistically produce pain hypersensitivity. Finally, although we reported that the number of circulating immune cells was not affected by the injury, we admit that the characterization on functional phenotypes of each subgroup had been omitted, for which potential changes could not be excluded.

## 5. Conclusion

Our study revealed that nerve injury like SNI could generate a long-lasting disturbance of systemic homeostasis, which was sexually dimorphic. Nerve injury did not affect overall immune cell numbers or proportions in both male and female mice, but it triggered a regulation of both pro- and anti-inflammatory cytokine and chemokine release in the serum, predominantly in male mice. Serum from both male and female mice with SNI contains likely different substances, but both induced pain hypersensitivity when transferred in a same-sex or cross-sex manner to naïve mice.

## Author contributions

WBSZ: Conceptualization, Investigation, Data curation, Formal analysis; Writing-original draft, review & editing.

XQS: Investigation, Data curation, Formal analysis, APZ: Data curation, Formal analysis,

MM: Conceptualization, Investigation, Writing-review & editing JM: Conceptualization, Writing-review & editing

JZ: Conceptualization, Project administration, Supervision, Funding acquisition, Writing-original draft, review & editing.

## Acknowledgements

The authors would like to thank Marija Landekic for help with flow cytometry experiment and analysis. We would also like to thank the McGill Flow Cytometry and Cell Sorting Facility for providing equipment and assistance for the flow cytometry experiments; the facility’s infrastructure is supported by the Canada Foundation for Innovation (CFI).

