## Supplementary figures for "The impact of nerve injury on the immune system across the lifespan is sexually dimorphic"

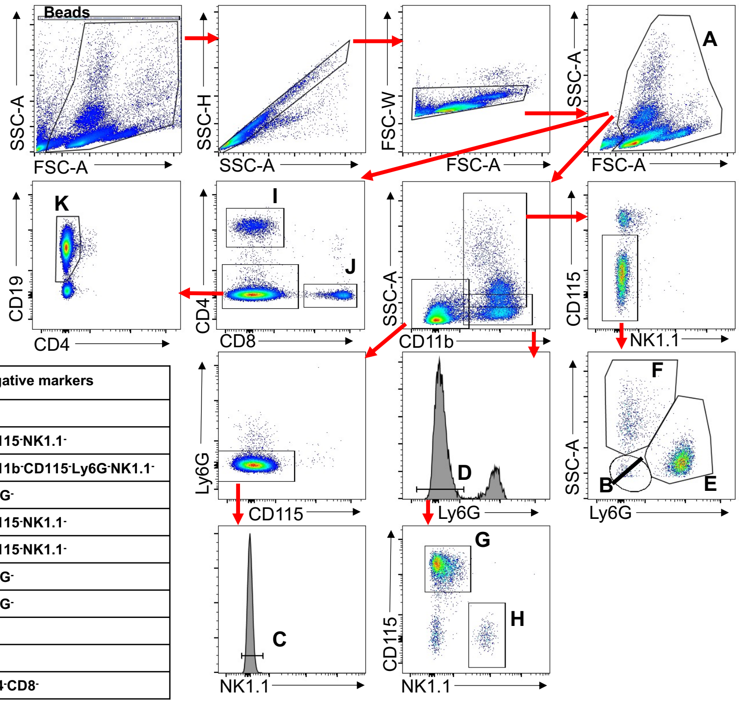


| | Population name | Panel | Positive markers | Negative markers |
|---|---|---|---|---|
| A | Total Leukocytes | | - | - |
| B | Granulocytes | 1 | CD11b$^+$ | CD115$^-$NK1.1$^-$ |
| C | Lymphocytes | 1 | | CD11b$^-$CD115$^-$Ly6G$^-$NK1.1$^-$ |
| D | Myeloid | 1 | CD11b$^+$ | Ly6G$^-$ |
| E | Neutrophils | 1 | CD11b$^+$Ly6G$^+$ | CD115$^-$NK1.1$^-$ |
| F | Other Granulocytes | 1 | CD11b$^+$ | CD115$^-$NK1.1$^-$ |
| G | Monocytes | 1 | CD11b$^+$CD115$^+$ | Ly6G$^-$ |
| H | NK Cells | 1 | CD11b$^+$NK1.1$^+$ | Ly6G$^-$ |
| I | CD4 T cells | 2 | CD4$^+$ | |
| J | CD8 T cells | 2 | CD8$^+$ | |
| K | B cells | 2 | CD19$^+$ | CD4$^-$CD8$^-$ |

**Figure S1. Gating strategy for circulating immune cells.** Representative gating strategy for all immune cell populations. Populations indicated with letters A-K next to the gates are identified in the adjacent table, along with positive and negative markers. The final gate for population B is an exclusion gate, where all events outside of the indicated gate are included in the population.

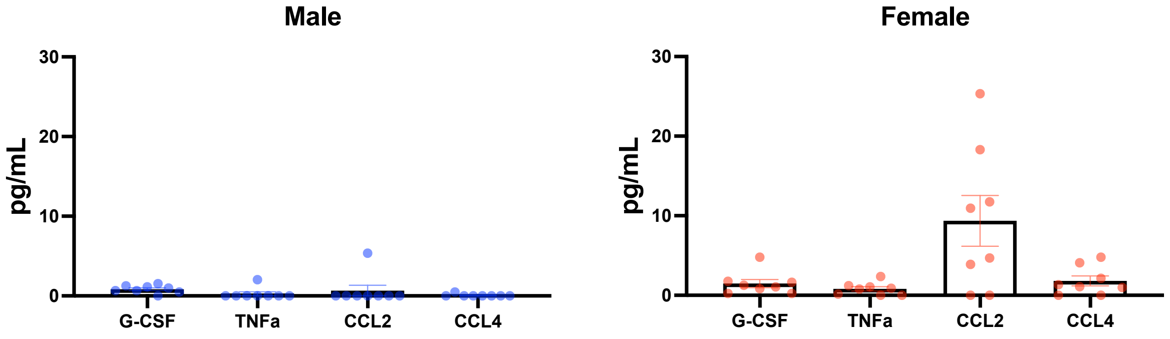


**Figure S2. Validation of Proteome Profiler results by Luminex assessment of serum cytokine concentrations in naive mice.** Individual serum samples taken from 2-3 months old naïve mice were assessed with a Luminex assay for cytokine concentrations of the targets G-CSF, TNFα, CCL2, and CCL4. All data are presented as the mean ± *SEM*. *n* = 8 mice/group

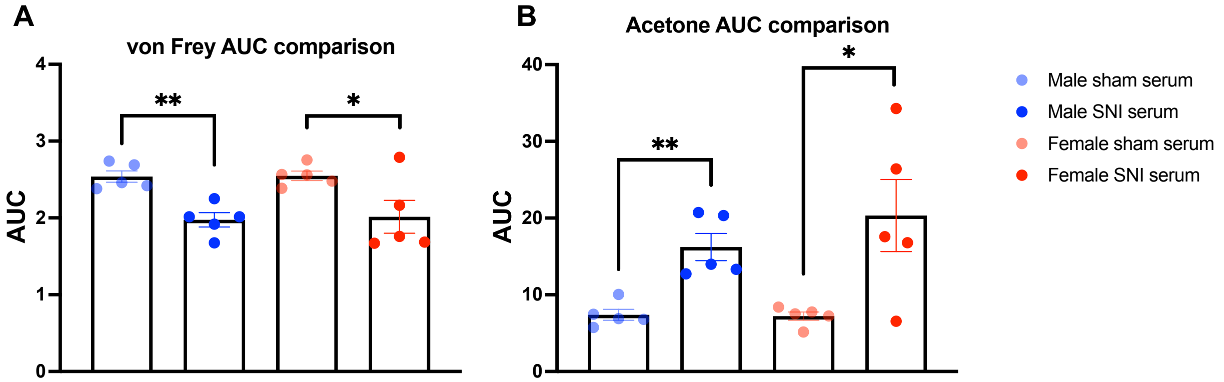


**Figure S3. Serum transfer pain behavior from day 1 – 3 analyzed with area under the curve.** Comparison of area under the curve with **(A)** von Frey and **(B)** acetone tests for all serum transfer experiments. All data are presented as the mean ± *SEM*. Significance was determined by unpaired t-test, *$P$<0.05, **$P$<0.01.
